# Investigating mechanisms underlying genetic resistance to Salmon Rickettsial Syndrome in Atlantic salmon using RNA sequencing

**DOI:** 10.1101/2020.12.03.410464

**Authors:** Carolina P. Moraleda, Diego Robledo, Alejandro P. Gutiérrez, Jorge del-Pozo, José M. Yáñez, Ross D. Houston

## Abstract

**Background:** Salmon Rickettsial Syndrome (SRS), caused by *Piscirickettsia salmonis,* is one of the primary causes of morbidity and mortality in Atlantic salmon aquaculture, particularly in Chile. Host resistance is a heritable trait, and functional genomic studies have highlighted genes and pathways important in the response of salmon to the bacteria. However, the functional mechanisms underpinning genetic resistance are not yet well understood. In the current study, a large population of salmon pre-smolts were challenged with *P. salmonis*, with mortality levels recorded and samples taken for genotyping. In parallel, head kidney and liver samples were taken from animals of the same population with high and low genomic breeding values for resistance, and used for RNA-Sequencing to compare their transcriptome profile both pre and post infection.

**Results:** A significant and moderate heritability (h^2^ = 0.43) was shown for the trait of binary survival. Genome-wide association analyses using 38K imputed SNP genotypes across 2,251 animals highlighted that resistance is a polygenic trait. Several thousand genes were identified as differentially expressed between controls and infected samples, and enriched pathways related to the host immune response were highlighted. In addition, several networks with significant correlation with SRS resistance breeding values were identified, suggesting their involvement in mediating genetic resistance. These included apoptosis, cytoskeletal organisation, and the inflammasome.

**Conclusions:** While resistance to SRS is a polygenic trait, this study has highlighted several relevant networks and genes that are likely to play a role in mediating genetic resistance. These genes may be future targets for functional studies, including genome editing, to further elucidate their role underpinning genetic variation in host resistance.

## BACKGROUND

Finfish aquaculture is a fast-growing industry with a worldwide production of 54.3 million tonnes during 2018, corresponding to an estimated value of USD 139.7 billion [1]. Atlantic salmon (*Salmo salar*) comprises 4.5% of global finfish trade, and demand for salmon has grown steadily since 2010 [1]. However, the expansion of salmon aquaculture has been associated with a concurrent increase in the occurrence and impact of infectious diseases, which can cause major welfare and production challenges. One of the most serious of these diseases is Salmon Rickettsial Syndrome (SRS), caused by the Gram-negative bacterium *Piscirickettsia salmonis*, which can cause severe morbidity and mortality in salmonid species. SRS is particularly problematic for salmon aquaculture in Chile, the world’s second largest producer, and is responsible for 47.5% of the total mortality due to infectious disease in this industry [2]. SRS has also been reported in other salmon-producing countries such as Norway, Ireland, Canada or Scotland. The morbidity and mortality caused by SRS occur at the seawater stage, where economic losses in relation to biomass are highest. The direct losses through mortality are exacerbated by indirect losses through reduced growth rates and premature harvests [3]. Several strategies for SRS control have been developed, such as vaccination, antibiotics and biosecurity measures, however, they have shown only partial efficacy under field conditions [3]. Development of novel strategies to control SRS requires improved knowledge of the genetic and functional aspects of *P. salmonis* host-pathogen interaction, such as the process of entry into host cells, intracellular replication, virulence mechanisms, and genetic variation in host response [3].

A promising avenue to mitigate the impact of SRS in Atlantic salmon aquaculture is to improve SRS disease resistance traits through selective breeding. This is possible due to naturally occurring genetic variation (heritability) for disease resistance, which has been observed in other infectious diseases impacting farmed populations of farmed salmonids [4–6]. Significant additive genetic variation for resistance to SRS has been found in various farmed populations, with family mortality levels ranging from 5% to 82% and heritability estimates from 0.11 to 0.41 [7, 8]. The genetic architecture of resistance to SRS has been studied using genome-wide association studies (GWAS) in populations of different salmonid species, suggesting that SRS resistance is a polygenic trait [9–11]. For such traits, genomic selection has been shown to be effective in increasing accuracy of breeding value prediction in commercial aquaculture breeding programmes [12, 13]. In the case of SRS resistance, the use of genomic information was shown to improve prediction accuracy by up to 30% compared to pedigree approaches [14].

While selective breeding and genomic selection for improved resistance to SRS can be performed without knowledge of the mechanisms underlying genetic resistance, understanding these mechanisms is a major goal for aquaculture research. Such information can yield novel disease treatment and mitigation options, including possible targets for vaccination and therapeutants. Furthermore, knowledge of functional genes and polymorphisms can be applied in functionally-enriched genomic selection, which can further improve prediction accuracy relative to the use of anonymous markers [15]. Finally, putative causative genes and variants can be targeted by CRISPR/Cas genome editing, initially to confirm their role, and ultimately to edit broodstock to carry resistant variants pending a suitable regulatory environment [16].

*P. salmonis* infects and replicated in salmonid macrophages, and stimulates a significant innate immune response together with an oxidative defence response [17, 18]. The host response to infection in Atlantic salmon has been assessed in a number of studies using microarrays and RNA-Sequencing. Their findings suggest that *P. salmonis* modulates the pro-inflammatory cytokine response, the iron deprivation system and the cytoskeletal reorganization, and interferes with protein transportation and vesicle trafficking to evade immune response, increase persistence and aid replication [19, 20]. However, while gene expression differences between families with different levels of resistance have been examined using microarrays [20], the functional mechanisms underpinning genetic variation in resistance to SRS remain poorly understood.

Therefore, the aims of this study were i) to evaluate the genetic architecture of SRS resistance in a large Atlantic salmon population from a commercial breeding programme, ii) to improve our understanding of the molecular basis of host response, and iii) to discover functional genes and pathways contributing to host genetic resistance to SRS.

## RESULTS

### Genetics of resistance to SRS

A large-scale *P. salmonis* injection challenge was performed on a population of salmon pre-smolts from a commercial breeding programme with fish distributed evenly across three tanks. The challenge was terminated after 47 days, and there were a total of 756 mortalities and 1509 survivors, corresponding to a mortality rate of 33%. The challenged fish started to die 17 days post-challenge, and mortality rate was consistent across the three tanks (Figure 1A). The estimated heritability of mortality as measured on the binary scale was 0.43 ± 0.04.

**Figure 1.**
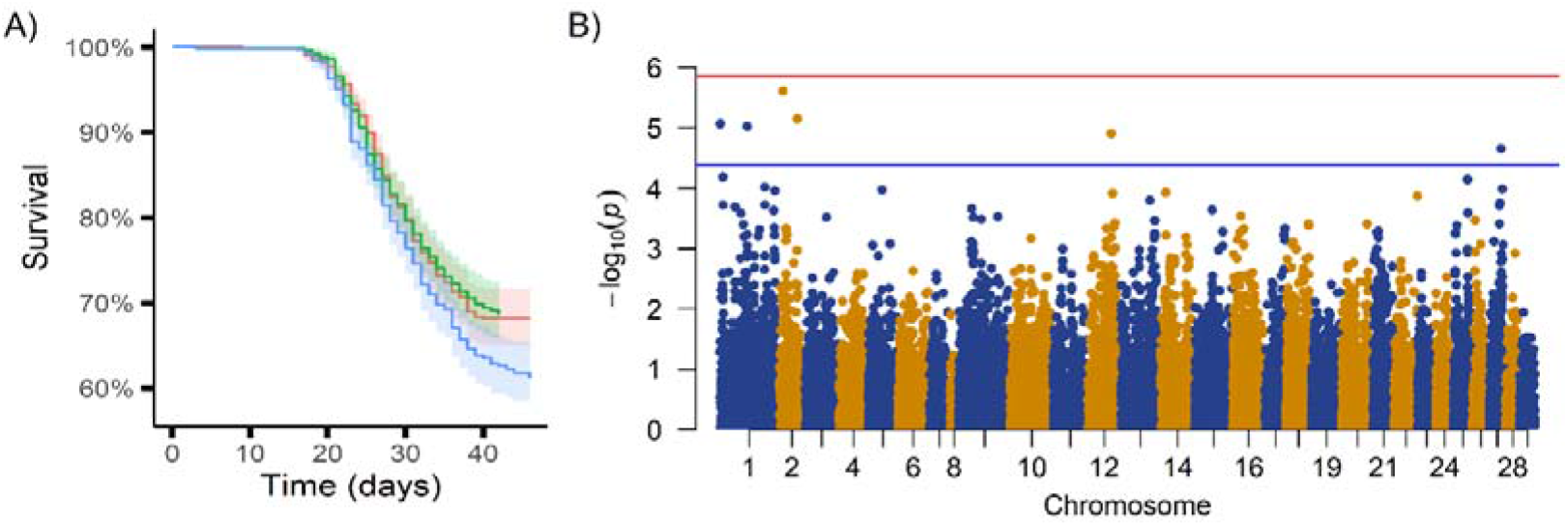
SRS disease challenge survival data and genome-wide association analysis. A) Percentage of survival in the population throughout the duration of the challenge in each of 3 tanks, and B) Manhattan plot showing the p-values of the GWAS for each SNP, the red line represents the Bonferroni corrected significance threshold and the blue line the suggestive significance threshold.

The genome-wide association analysis revealed a polygenic architecture for the trait of resistance to SRS, although a few SNPs reached the suggestive level of significance (Figure 1B). These SNPs were situated on chromosomes 1, 2, 12 and 27, indicative of putative QTL on these chromosomes. However, no single SNP explained more than 1% of the genetic variation in resistance to SRS.

### Transcriptomic response to SRS infection

To examine the transcriptomic response to infection, 48 fish were euthanized and sampled pre-challenge, 3 days post-challenge and 9 days post-challenge from the same tank (total n = 144). Head kidney and liver samples were obtained from each animal and stored in RNAlater at 4 °C for 24 h, and then at −20°C until RNA extraction. A total of 133 samples were then selected for RNA sequencing (74 liver and 59 head kidney samples; Supplementary file 2) based on (i) high and low EBVs for resistance to SRS, and (ii) RNA quality. An average of ~40M reads per sample were produced using RNA Sequencing of the head kidney and liver samples collected at 3 and 9 days post-challenge. Hierarchical clustering of all the samples using gene expression data clustered head kidney and liver separately, as expected (Figure 2A). Principal Component Analysis was performed in each tissue separately to assess the sample clustering within tissue. Liver samples showed a clear separation between controls and the 9 days post infection samples, with the samples from 3 days post infection falling in between and showing a significant overlap with the other two groups (Figure 2B). In the case of head kidney, the infected samples clustered separately from controls, but a clear separation between 3 and 9 days post infections was not observed (Figure 2C).

**Figure 2.**
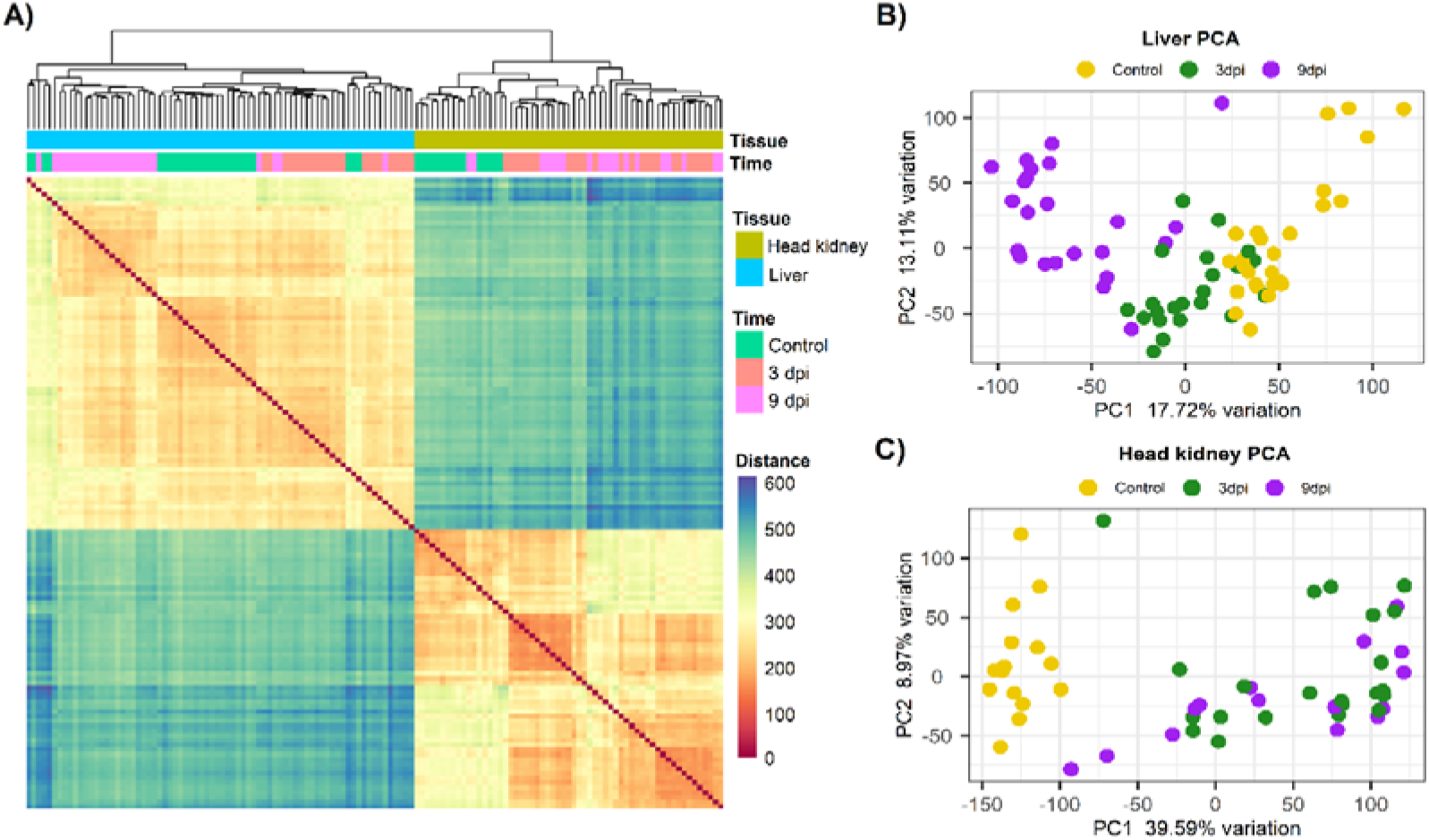
Sample clustering based on RNA-Sequencing data from liver and head kidney samples. A) Hierarchical clustering of all samples, and B) principal component analyses of the liver samples and C) of the head kidney samples.

Differential expression analyses between controls and infected samples highlighted a very large number of differentially expressed genes (9K to 12.5K per comparison, FDR p-value < 0.05), which was expected considering the high statistical power associated with the large sample size in this experiment. To facilitate downstream analyses and interpretation, only genes with FDR p-value < 0.01, normalized mean expression > 10 reads, and log2FC > 0.5 were considered. This resulted in 5,000 to 7,000 differentially expressed genes in each comparison, with an even number of up-regulated and down-regulated genes (Table 1, Figure 3, Supplementary file 3). Several innate immune genes were regulated in response to SRS, including interleukins, tumor necrosis factor related genes, caspases and interferon genes (Figure 3).

**Figure 3.**
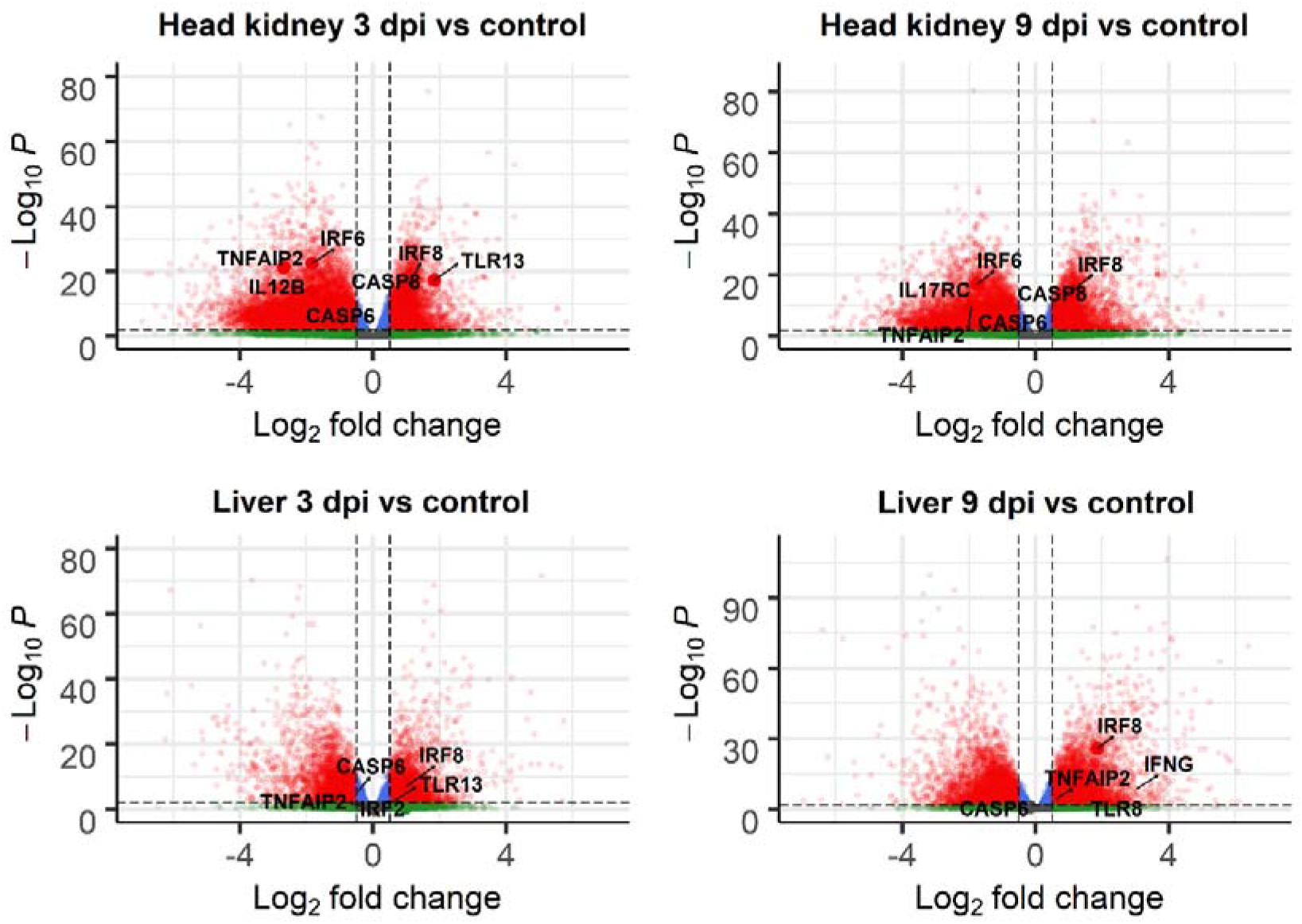
Volcano plots of RNA-Seq data comparing control vs SRS infected samples. Each point in the plots represents a gene, with its log2 fold change in the x-axis and its −log10 p-value in the y-axis. Positive fold change means upregulated in infected samples. Genes are classified in 4 categories depending on their FC and FDR corrected p-value: i) grey = p-value > 0.01 and log2 fold change between −0.5 and 0.5; ii) green = p-value > 0.01 and log2 fold change < −0.5 or > 0.5; iii) blue = p-value < 0.01 and log2 fold change between −0.5 and 0.5; and iv) red = p-value < 0.01 and log2 fold change < −0.5 or > 0.5).

**Table 1.** Differential expression between control and SRS challenged samples. Upregulated and downregulated refers to the challenged samples versus the unchallenged controls.

|  | 3 dpi |  | 9 dpi |  |
| --- | --- | --- | --- | --- |
|  | Upregulated | Downregulated | Upregulated | Downregulated |
| **Head kidney** | 2603 | 2831 | 2705 | 2529 |
| **Liver** | 3117 | 3453 | 3260 | 3111 |

Between 15 and 55 KEGG pathways were enriched for differentially expressed genes in the four comparisons (Figure 4, Supplementary File 4). Generally, immune pathways such as cytokine-cytokine receptor interactions, apoptosis, and Toll-like receptor signaling showed enrichment for gene upregulation in both organs, albeit more strongly in head kidney than liver at 3dpi. TNF signaling and bacterial invasion of epithelial cells were only enriched for upregulated genes in head kidney, while evidence for *Staphylococcus aureus* infection and phagosome upregulation was liver-specific. Energy metabolism pathways showed evidence for downregulation in both organs, including glycolysis / gluconeogenesis or fatty acid degradation (Figure 4).

**Figure 4.**
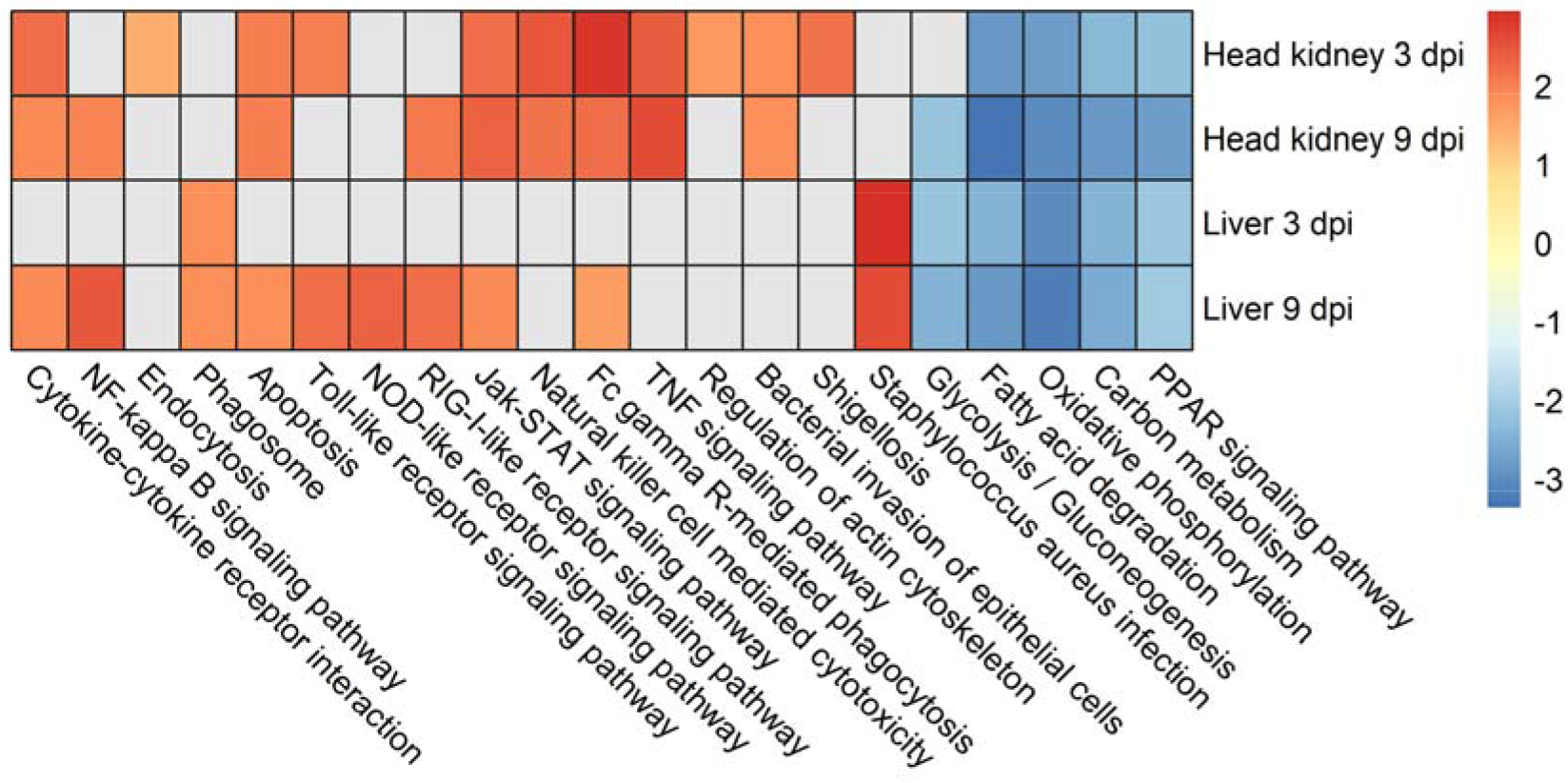
KEGG pathways enriched for genes showing significant differential expression between SRS infected and control samples. Heatmap showing the fold enrichment of selected KEGG pathways showing significant up- (positive values) or down-regulation (negative) in response to SRS infection.

### Signatures of resistance to SRS

SRS resistance breeding values for all the RNA-Seq animals were estimated according to the linear mixed model described in the methods. To investigate the association between gene expression and resistance to SRS, a network correlation analysis was performed. Head kidney and liver transcriptomes clustered into 30 and 22 putative gene networks respectively, with each network containing between 25 and 7,000 genes. The correlation between the SRS resistance EBVs at each time point and average network gene expression (Supplementary Figure 1) revealed significant associations for 6 and 2 gene networks in head kidney and liver, respectively (|r| > 0.45, p < 0.001; Supplementary files 5 and 6, respectively), suggesting that these networks may play a functional role in defining host resistance to SRS. KEGG enrichment analysis of the gene networks associated with resistance revealed genes involved in the apoptotic processes, such as BCL2L1, ITP3 and BNIP3, in the Cytoskeletal reorganization pathway such as SPTB, and in Bacterial invasion and Intracellular trafficking such as CBL and RAB9A (Figure 5).

**Figure 5.**
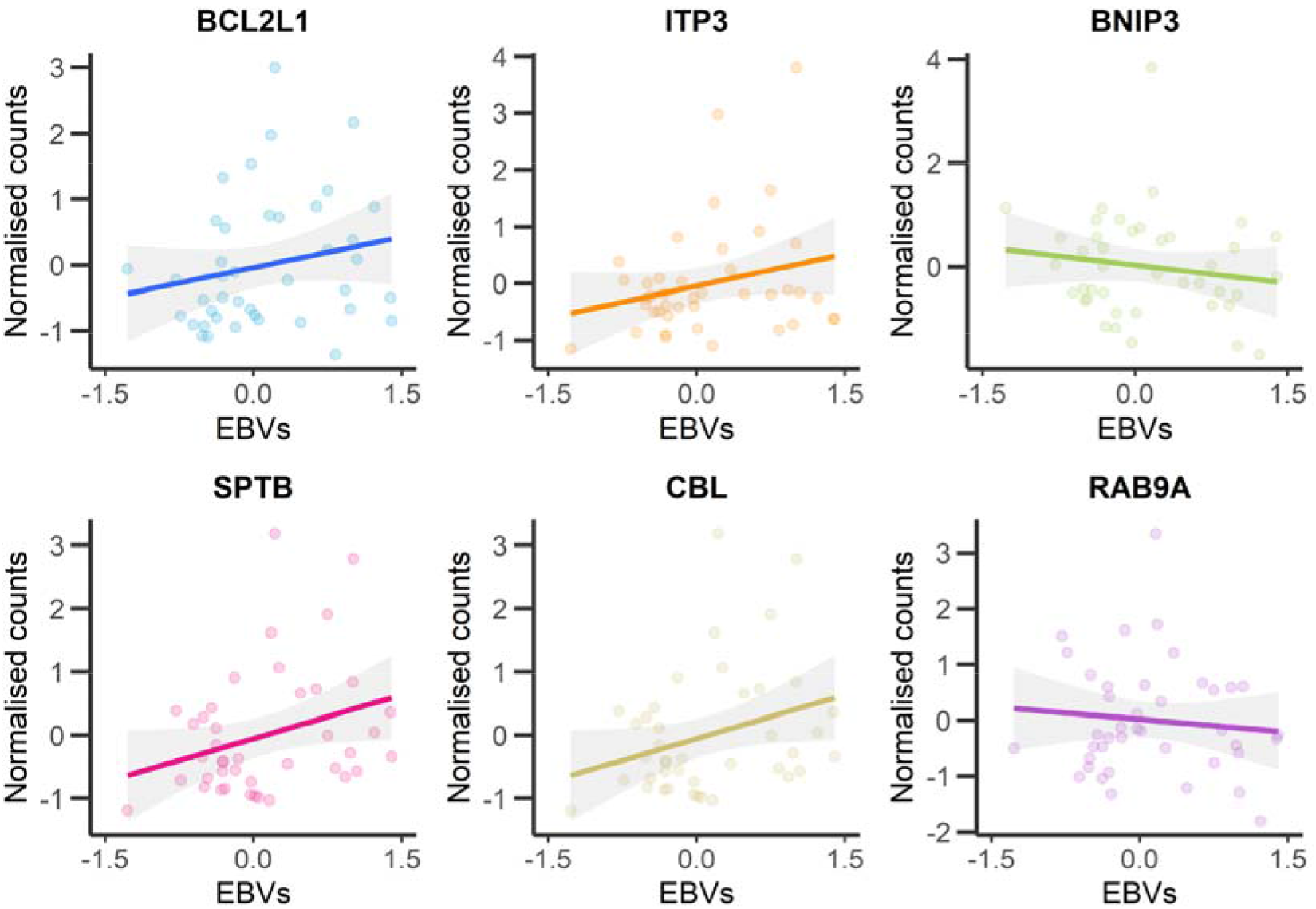
Correlation between gene expression and breeding values for resistance to SRS. Correlation between the expression of 6 genes of interest (normalized read counts) and the estimated breeding values (EBVs) for resistance to SRS. The six genes are Bcl-2-like protein 1 (BCL2L1), Ion transport peptide 3 (ITP3), BCL2/adenovirus E1B 19 kDa protein-interacting protein 3 (BNIP3), Spectrin beta chain (SPTB), E3 ubiquitin-protein ligase CBL (CBL), and Ras-related protein Rab-9a (RAB9A).

## DISCUSSION

Improving our understanding of the functional basis of genetic resistance and host response to SRS in Atlantic salmon is valuable for the development of new strategies of disease control. To this end, this large-scale study has provided further evidence for significant heritability of host resistance to SRS, and suggested that the genetic architecture of resistance is polygenic in nature. Furthermore, RNA-Sequencing of liver and head kidney samples from SRS-challenged salmon pre-smolts highlighted a large-scale up-regulation of immune pathways and down-regulation of energy metabolic pathways compared to controls.

Resistance to SRS in the population studied herein has a moderate level of genetic control, with a heritability estimate of 0.43 (binary survival). This estimate is towards the upper limit of those reported in previous studies for Atlantic salmon, which ranged between 0.11 and 0.41 [7, 21, 22], and is also similar to those reported for resistance to SRS in rainbow trout (ranging between 0.38 and 0.54) [8, 23], but somewhat higher to the values found in coho salmon (ranging between 0.16 to 0.31) [24, 25]. The genetic variation in resistance to SRS appears to be polygenic in nature, without any significant major QTL, and suggestive QTLs on only four chromosomes. This polygenic architecture was also reported in previous studies [9–11]. Chromosomes 1 and 12 have also been found harbouring genomic regions associated with resistance to SRS in previous studies carried out in a different Atlantic salmon population, raising the possibility the QTL are the same [9, 10]. The putative QTL found herein on chromosomes 2 and 27 identified here differ from previous studies, which can be explained by differences in disease challenge conditions (discussed below), different genetic background between populations and the polygenic nature of the trait. Nonetheless, the moderate heritability and polygenic architecture of resistance to SRS in Atlantic salmon make this trait an ideal candidate for genomic selection in salmon breeding programmes, which has proved to be an efficient method to select for resistance to SRS and other diseases with a polygenic background in salmon [14, 26–29]. However, it should be noted that the intraperitoneal injection model used for SRS challenges will have significant impact on the interpretation of the trait of genetic resistance. For example, the route of entry for *P. salmonis* is via epithelial tissues (skin and gills), and these tissues are known to present a critical barrier against bacterial infection [30]. The intraperitoneal injection bypasses this, and therefore it is to be expected that only part of the mechanisms of genetic resistance are being captured. For this reason, benchmarking genetic resistance measured in the laboratory injection challenge with mortality levels observed in the field is an important consideration [31].

SRS infected animals showed major transcriptional differences compared to uninfected controls in both the head kidney and the liver, involving the differential expression of thousands of genes, similarly to previous studies that also reported a significant gene expression modulation in liver and head kidney in response to SRS [20, 32, 33]. Several important innate immune response pathways were up-regulated in both organs, such as Apoptosis, NOD-like receptor signalling, NF-kappa B signalling and Bacterial invasion of epithelial cells (Figure 4). Likewise, several energy metabolism pathways are down-regulated in response to the infection, probably as a result of diversion of cellular resources towards immune response, as has been suggested in previous studies of macrophage cell lines response to *P. salmonis* infection [17]. The integration of the transcriptomic response to infection and the gene network analysis to identify signatures of resistance to SRS allowed us to identify four key biological processes that seem to be important for the outcome of the infection: i) cytoskeleton reorganization, ii) apoptosis, iii) bacterial invasion and intracellular trafficking, and iii) the inflammasome.

### Cytoskeleton reorganization

Genes and pathways related to cytoskeleton reorganization featured heavily in the lists of differential expression genes in response to infection. The cytoskeleton plays an active role in the innate immune response: cytoskeletal activation is involved in pathogen detection, phagocytosis, cell-cell signalling, cell migration, and secretion [34]. Furthermore, major disruptions in actin components have been described during the infection process of intracellular bacteria such as *Legionella pneumophila*, *Coxiella burnetii* and *Listeria monocytogenes* [35–38]. Similarly, *P. salmonis* modulates the cytoskeleton by inducing actin depolymerization [39], which results in cytoskeletal reorganization [19]. This is consistent with our results, where several cytoskeleton associated genes showed high correlation with estimated breeding values for resistance. A notable example is the Rho-associated coiled-coil kinase 1 (ROCK1; r = 0.27), a serine/threonine kinase downstream effector of the Rho family, described as an essential regulator of actin cytoskeleton [40]. ROCK kinases participate in the bacterial invasion of *Coxiella burnetii* in human cells, and the use of ROCK inhibitors during infection hampered the bacterial internalization process [41]. Furthermore, genes highly correlated with SRS susceptibility such as SPTB (r = −0.57) and SEPTIN3 (−0.42) are cytoskeleton constituents that participate in protein linking (SPTB; [42]) and GTP-binding (SEPTIN3; [43]), respectively. This high correlation of these genes with susceptibility may be explained by the availability of actin in these structures, which is a target for modulation by the bacterium during cytoskeletal depolymerisation, and therefore disrupting this modulation of the cytoskeleton may be a strategy to increase resistance to SRS.

### Apoptosis and cell survival promotion

Apoptosis is a programed cell-death mechanism essential to development and maintenance of homeostasis [44]; but induction of apoptosis has also been observed during bacterial and viral infection, hampering microbial replication and dissemination [45]. Intracellular bacteria actively modulate cellular apoptosis to enable their replication within the cells [46]. Previous studies suggest that *P. salmonis* modulates the apoptotic process of the host as a strategy to ensure intracellular survival [19, 47]. In line with this, apoptotic genes and pathways were heavily modulated during SRS infection in the current study. Furthermore, the expression of two different inhibitors of apoptosis, BCL2L1 (r = −0.62) and ITP3 (r = −0.50), was negatively correlated with resistance to SRS. BCL2L1 inhibits caspase-1 activation by interfering with NLRP1 oligomerization, a key component of the inflammasome immune response [48], and ITP3 has an anti-apoptotic effect in mammalians cancer cells [49]. In contrast, apoptosis promoting genes, such as BNIP3 (r = 0.33) [50, 51] and Bim (BCL2L11 r = 0.18) [52], were positively correlated with genetic resistance. These findings support the hypothesis that apoptosis is initiated as a host strategy to mitigate pathogen dissemination, which is subverted by SRS to promote cell survival and bacterial replication.

### Bacterial invasion and intracellular trafficking

The intracellular environment provides diverse advantages to pathogens, for example protection against humoral and complement-mediated host defence mechanisms, and availability of nutrients and direct access to metabolic pathways to modulate in their favour. In order to stablish an intracellular infection, pathogens utilise a wide range of mechanisms for internalization and survival [53]. Once inside host cells, *P. salmonis* is capable of establishing intracellular infections, and replicate in macrophages within cytoplasmic vacuole-like structures [54]. In *P. salmonis*, this is facilitated by a virulence factor that encodes a type IVB secretion system [17, 55]. The Dot/Icm type IVB secretion system allows bacteria to translocate proteins into host cells, and manipulate host pathways [56]. In *P. salmonis*, this may involve modulation of the host cell intracellular trafficking, leading to disrupted phagosome-lysosome pathogen clearance [55]. Interestingly, in this study key genes participating in intracellular trafficking such as RAB1B (r = 0.24) and RAB9A (r = 0.63) are positively correlated with genetic resistance to SRS. RAB1B is a Rab protein modulated by *Legionella pneumophila* Dot/Icm T4SS effectors to recruit endoplasmic reticulum-derived vesicles to stablish bacterial replication vacuoles [57]. Conversely, RAB9A is involved in the transport between endosome vesicles and the trans Golgi network [58], and is interrupted by *Salmonella enterica* SifA effector to attenuate the lysosomal activity in *Salmonella* containing vacuoles (SCV) [59]. In the current study a strong negative correlation was found between the gene CBL (r = −0.52) and resistance to SRS, suggesting that *P. salmonis* virulence factors may target this gene to facilitate bacterial internalization. Furthermore, E3 ubiquitin-protein ligase CBL-like isoform X1 (CBL) was found in chromosome 2, located in the most significant QTL region for resistance to *P. salmonis* infection. Interestingly, *Listeria monocytogenes*, another intracellular bacteria, expresses surface proteins to modulate host proteins like Met and CBL and hijack the clathrin-dependent endocytosis process [60], and previous studies indicate that *P. salmonis* internalization process is mediated by clathrin endocytosis [39].

### Inflammasome

Another interesting result was the large number of genes differentially expressed in response to infection involved in the inflammasome. The inflammasome is an intracellular sensing system activated by a broad range of microorganisms that has a pivotal role in the innate immune response to infection [61]. Activation of the inflammasome initiates a signalling cascade that culminates in caspase-1 expression and maturation of the proinflammatory cytokine IL-1β [62]. Numerous studies suggest that genes participating in the inflammasome assembly may be conserved in teleost fish [63, 64]. Moreover, gene activation of inflammasome associated components such as NLRP1, ASC and caspase-1 has been described in response to bacterial infection in zebrafish (*Danio rerio*) and turbot (*Scophthalmus maximus*) [65, 66]. In the current study, genes involved in the activation of the inflammasome had higher expression on average in resistant fish, suggesting that overexpression of this pathway could be protective during SRS infection. The expression of NLRP1, a sensor that initiates the inflammasome response, is significantly positively correlated with genetic resistance (r = 0.20). NLRP1 is a NOD-like receptor (NLR) that detects pathogen molecules and triggers the activation of effector caspases (caspases 1, 4, 5 and 11) [65]. Similarly, NLRC3 is another component of the inflammasome positively correlated with resistance (r = 0.31). While in humans it has been described as an inhibitor of the innate immune response through the inhibition of NF-kB activity [64], in teleosts NLRC3 expression is significantly increased in mucosal tissue after exposure to bacteria, implying an involvement in the early immune response [67, 68]. In contrast, NLRP12 (r = −0.4774) is a regulator of inflammation which acts as a suppressor of pro-inflammatory cytokines interfering with the NF-kB pathway [69], and therefore its negative correlation with genetic resistance suggests that the activation of the inflammasome pathway is beneficial in response to SRS. In summary, these findings suggest that the activation of the inflammasome pathway is important for a successful immune response against *P.salmonis*.

## CONCLUSIONS

This study highlights a significant genetic component to SRS resistance in Atlantic salmon, underpinned by a polygenic architecture. The RNA-Sequencing comparison of control and infected fish identified a major signature of host response evident in both head kidney and liver tissues. When comparing this response between individual fish of high and low resistance breeding values, several interesting gene expression networks were identified that correlate with genetic resistance. These include genes related to cytoskeleton, apoptosis and cell survival, bacterial invasion/intracellular trafficking, and the inflammasome. Considering the scale and complexity of the transcriptomic response observed in salmon challenged with *P. salmonis*, and the lack of any significant QTL associated with host resistance, the potential mechanisms leading to genetic resistance are likely to be heterogeneous and vary across different families and individuals. However, the pathways and genes highlighted by this study are potential candidates for functional studies, and downstream applications in salmon production. For example, strategies to increase resistance to the bacteria can focus on disrupting its modulation of cellular homeostasis (i.e. cytoskeleton or apoptosis) or on boosting the immune processes that prevent or restrain the infection (i.e. inflammasome). Such strategies may include CRISPR/Cas knockout or modulation in cell line models, or ultimately *in vivo* to interrogate the impact of perturbation of the identified genes on genetic resistance.

## MATERIALS AND METHODS

### Experimental design

2,265 Atlantic salmon pre-smolts (average weight 174 g) from 96 full sibling families from the breeding population of AquaInnovo (Salmones Chaicas, Xth Region, Chile) were experimentally challenged with *Piscirickettsia salmonis* (strain LF-89) in 3 × 7 m^3^ tanks. Fish were intraperitoneally injected with 0.2 mL of a 1/2030 dilution of *P. salmonis*. This dose was expected to cause a population-level mortality of close to 50%, based on a pre-challenge of 300 fish from the same families challenged with different doses of the bacteria. The main challenge was terminated after 47 days, after mortality levels had dropped to close to baseline levels. Caudal fin clips were taken from all mortality and survivor fish for future DNA extraction and genotyping.

For RNA sequencing, 48 fish were sampled pre-challenge, 3 days post-challenge and 9 days post-challenge from the same tank, for a total of 144 fish. Head kidney and liver samples were obtained from each animal and stored in RNAlater at 4 °C for 24 h, and then at −20°C until RNA extraction.

### Genotyping and imputation

DNA was extracted from the fin clips of the challenged fish using a commercial kit (Wizard Genomic DNA Purification Kit, Promega), following the manufacturer’s instructions. All samples where genotyped with a panel of 968 SNPs (Supplementary file 1) chosen as a subset of the SNPs from a medium density SNP array [70] using Kompetitive Allele Specific PCR (KASP) assays (LGC Ltd, UK). A population containing full-siblings of the challenged animals had previously been genotyped with a SNP panel of 45,818 SNPs (n = 1,056, [70]; Supplementary file 1), and the experimental population was imputed to ~46K SNPs using FImpute v.2.2 [71]. Imputation accuracy was estimated by 10-fold cross validation, masking all SNPs except the 968 SNP panel for 10% of the 1,056 genotyped full-sibs, and then assessing the correlation between the true genotypes and the imputed genotypes for the remainder of the SNPs. All imputed SNPs showing imputation accuracy below 80% were discarded. The average imputation accuracy for the 39,416 SNPs retained (Supplementary file 1) was of 95%. Further details about the low-density SNP panel and imputation methods can be found in Robledo et al. (2019)[72]. The imputed genotypes were then filtered and removed according to the following criteria: SNP call-rate < 0.9, individual call-rate < 0.9, FDR rate for high individual heterozygosity < 0.05, identity-by-state > 0.95 (both individuals removed), Hardy-Weinberg equilibrium p-value < 10^−6^, minor allele frequency < 0.01. After filtering 38,028 markers and 2,345 fish remained for the downstream analyses.

### Estimation of genetic parameters

The phenotype of resistance to SRS was measured as binary survival, recording mortalities as 0 and survivors as 1. Genetic parameters for SRS resistance were estimated using the genomic relationship matrix (**G**-matrix) to model the additive genetic relationship between animals in ASReml 4.1 [73] using he following linear mixed model:

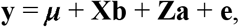

where **y** is a vector of observed phenotypes, μ is the overall mean of phenotype records, **b** is the vector of fixed effects which includes tank as factor and weight at the start of the challenge as covariate, **a** is a vector of additive genetic effects distributed as ~N(0,**G**σ^2^a) where σ^2^a is the additive (genetic) variance, **G** is the genomic relationship matrix. **X** and **Z** are the corresponding incidence matrices for fixed and additive effects, respectively, and **e** is a vector of residuals. The identity-by-state genomic relationship matrix (**G**) was calculated using the GenABEL R package (“gkins” function; [74]) kinship matrix [75], multiplied by two and inverted.

### Single-SNP genome-wide association study

The single-SNP GWAS was performed using the GenABEL R package [74] by applying the mmscore function [76], which accounts for the relatedness between individuals applied through the GenABEL [74] genomic kinship matrix [75]. Significance thresholds were calculated using a Bonferroni correction where genome-wide significance was defined as 0.05 divided by number of SNPs [77] and suggestive as one false positive per genome scan (1 / number SNPs).

### RNA extraction and sequencing

For all the 288 head kidney and liver samples, a standard TRI Reagent RNA extraction protocol was followed. Briefly, approximately 50 mg of tissue was homogenized in 1 ml of TRI Reagent (Sigma, St. Louis, MO) by shaking using 1.4 mm silica beads, then 100 μl of 1-bromo-3-chloropropane (BCP) was added for phase separation. This was followed by precipitation with 500 μl of isopropanol and posterior washes with 65-75 % ethanol. The RNA was then resuspended in RNAse-free water and treated with Turbo DNAse (Ambion). Samples were then cleaned up using Qiagen RNeasy Mini kit columns and their integrity was checked on Agilent 2200 Bioanalyzer (Agilent Technologies, USA). A total of 133 samples were selected for RNA sequencing (74 liver and 59 head kidney samples; Supplementary file 2) based on their EBVs for resistance to SRS and RNA quality. Thereafter, the Illumina Truseq mRNA stranded RNA-Seq Library Prep Kit protocol was followed directly. Libraries were checked for quality and quantified using the Bioanalyzer 2100 (Agilent), before being sequenced on 16 lanes of the Illumina Hiseq 4000 instrument using 75 base paired-end sequencing at Edinburgh Genomics, UK. Raw reads have been deposited in NCBI’s Sequence Read Archive (SRA) under BioProject accession number PRJNA669807.

### Read mapping

The quality of the sequencing output was assessed using FastQC v.0.11.5 (http://www.bioinformatics.babraham.ac.uk/projects/fastqc/). Quality filtering and removal of residual adaptor sequences was conducted on read pairs using Trimmomatic v.0.38 [78]. Specifically, Illumina specific adaptors were clipped from the reads, leading and trailing bases with a Phred score less than 20 were removed and the read trimmed if the sliding window average Phred score over four bases was less than 20. Only reads where both pairs were longer than 36 bp post-filtering were retained. Trimmed reads were then pseudoaligned against the Atlantic salmon reference transcriptome (ICSASG_v2 Annotation Release 100; [79]) using kallisto v0.44.0 [80].

### Differential expression

Transcript level expression was imported into R v3.6 [81] and summarised to the gene level using the R/tximport v1.10.1 [82]. Gene count data were used to estimate differential gene expression using the Bioconductor package DESeq2 v.3.4 [83]. Briefly, size factors were calculated for each sample using the ‘median of ratios’ method and count data was normalized to account for differences in library depth. Next, gene-wise dispersion estimates were fitted to the mean intensity using a parametric model and reduced towards the expected dispersion values. Finally a negative binomial model was fitted for each gene and the significance of the coefficients was assessed using the Wald test. The Benjamini-Hochberg false discovery rate (FDR) multiple test correction was applied, and transcripts with FDR < 0.01, normalized mean read counts > 10 and absolute log2 fold change values (FC) > 0.5 were considered differentially expressed genes. Hierarchical clustering and principal component analyses were performed to visually identify outlier samples, which were then removed from the analyses. The R packages “pheatmap”, “PCAtools” and “EnhancedVolcano” were used to plot heatmaps, PCAs and volcano plots, respectively. Kyoto Encyclopedia of Genes and Genomes (KEGG) enrichment analyses were carried out using KOBAS v3.0.3 [84]. Briefly, salmon genes were annotated against KEGG protein database [85] to determine KEGG Orthology (KO). KEGG enrichment for differentially expressed gene lists was tested by comparison to the whole set of expressed genes in the corresponding tissue using Fisher’s Exact Test (genes with mean normalized count values > 10). KEGG pathways with ≥ 5 DE genes assigned and showing a Benjamini-Hochberg FDR corrected p-value < 0.05 were considered enriched for differential expression.

### Network correlation analysis

Network correlation analyses were performed in R v3.6 [81] using the WGCNA package v1.69 [86]. Read counts after variance stabilizing transformation in DESeq2 [83] were used as measure of gene expression. Co-expression networks were then built using a power of 10, and clusters of genes were grouped into different color modules, allowing a minimum of 25 genes per module. Correlation between network summary profiles and external traits was quantified, and network trait associations showing |r| > 0.45 and p < 0.001 were considered significant. Thereafter, Kegg enrichment analyses were performed for the significantly associated networks using KOBAS 3.0.3 [84] as described above.

## DATA AVAILABILITY

RNA sequencing raw reads have been deposited in NCBI’s Sequence Read Archive (SRA) under BioProject accession number PRJNA669807.

## ETHICS DECLARATIONS

The challenge experiments were performed under local and national regulatory systems and were approved by the Animal Bioethics Committee (ABC) of the Faculty of Veterinary and Animal Sciences of the University of Chile (Santiago, Chile), Certificate N° 01-2016, which based its decision on the Council for International Organizations of Medical Sciences (CIOMS) standards, in accordance with the Chilean standard NCh-324-2011.

## CONSENT FOR PUBLICATION

Not applicable.

## AUTHOR CONTRIBUTIONS

RH and JY were responsible for the concept and design of this work. AG performed the molecular biology experiments. CM and DR performed data analysis and interpretation. CM, DR, RH, JP drafted the manuscript. All authors read and approved the final manuscript.

## ACKNOWLEDGEMENTS

We would like to acknowledge Benchmark Genetics Chile and Salmones Chaicas for providing the biological material and phenotypic records of the experimental challenges, and providing high-density genotypes of full-sibs of our experimental population for imputation. We would also like to acknowledge Edinburgh Genomics for performing the RNA Sequencing.

