## Supplementary figures and images for "Investigating mechanisms underlying genetic resistance to Salmon Rickettsial Syndrome in Atlantic salmon using RNA sequencing"

### Supplementary Figure 1

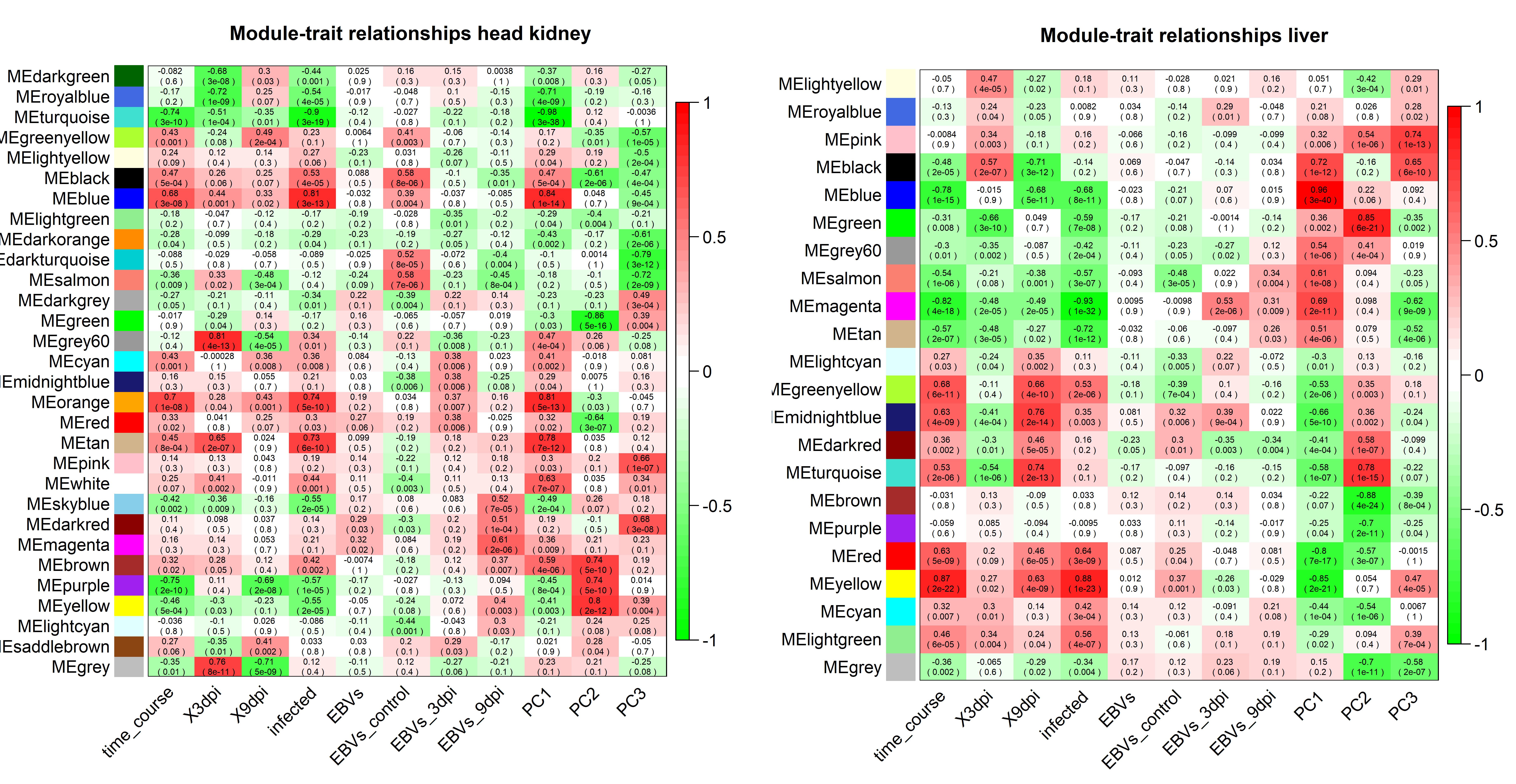
